# Pre-dispersal conditions and presence of opposite sex modulate density dependence and sex bias of dispersal

**DOI:** 10.1101/146605

**Authors:** Abhishek Mishra, Sudipta Tung, V.R. Shree Sruti, Mohammed Aamir Sadiq, Sahana Srivathsa, Sutirth Dey

**Author notes:** **Name and address of the corresponding author: Sutirth Dey** Associate Professor, Biology Division, Indian Institute of Science Education and Research, Pune, Dr. Homi Bhabha Road, Pune, Maharashtra, India 411 008, **Email:**.

## Abstract

Density-dependent dispersal (DDD) has been demonstrated in many species and has several ecological and evolutionary consequences. Yet we know little about how robust DDD is to the various conditions experienced by individuals. In this study, we use three independent experiments on laboratory populations of *Drosophila melanogaster* to examine the effects of pre-dispersal adult density, sex of the dispersers and presence of mates on the robustness of DDD patterns. We show that DDD can be greatly affected by both pre-dispersal density and interaction between the sexes. Moreover, the direction of sex-biased dispersal can reverse completely due to an interaction between the pre-dispersal and dispersal densities. We also show that interaction between the sexes can lead to negative DDD at the population level, even if, by themselves, neither sex exhibits DDD. Finally, we discuss potential implications of our results for processes like evolutionary rescue from extinctions and genetic divergence of populations.

## 1 INTRODUCTION

Dispersal is a complex process which is caused and modulated by a large number of factors such as landscape structure, resource availability, and individual phenotype (reviewed in Matthysen 2012). One of the factors that has been often implicated as a cause of variation in dispersal is population density (Matthysen 2005), and density-dependent dispersal (DDD) in turn has been shown to have major ecological and evolutionary implications. For instance, disease vectors can exhibit DDD (Trewhella *et al.* 1988; Gürtler *et al.* 2009), which could further modulate the rate and spread of diseases. Moreover, DDD is expected to have pronounced effects on the synchrony and dynamics of spatially structured populations (Amarasekare 2004; Lecomte *et al.* 2004), which is of particular significance in the context of endangered species (Rouquette & Thompson 2007). Furthermore, DDD is shown to have population genetic consequences (Aars & Ims 2000), indicating that it could play a role, along with other plastic dispersal behaviors, in macro-evolutionary processes such as speciation (Arendt 2015).

Not surprisingly therefore, density-dependence of dispersal has been thoroughly investigated both theoretically and empirically. Theoretical investigations have focused on the effects of DDD on population distribution (Namba 1980), synchrony (Liu *et al.* 2016), and source-sink dynamics (Amarasekare 2004), as well as the conditions under which DDD is expected to evolve (Travis *et al.* 1999; Poethke & Hovestadt 2002). Empirical studies across multiple taxa show that increased intraspecific competition is expected to drive higher emigration at high densities, leading to positive DDD (De Meester & Bonte 2010; Lutz *et al.* 2015). In contrast, some other organisms show a reduced dispersal with increasing population density, either due to a reduced availability of mates and resources (Lambin 1994), or social costs (Hestbeck 1982). In spite of this wealth of studies on DDD, an issue that remains relatively unexplored is the robustness of DDD patterns under different conditions.

In most studies on DDD, the current density (i.e., the density observed at the beginning of the dispersal phase) is taken into account while quantifying dispersal (e.g., Bitume *et al.* 2013; Schultz *et al.* 2017) thus neglecting the history of the population. However, dispersal can be affected by the physiology of the dispersers (Zera & Denno 1997), which in turn is expected to be a function of the pre-dispersal conditions. Therefore, intuitively speaking, it is possible that DDD patterns can themselves be modulated by the pre-dispersal conditions of the organisms. This line of reasoning is consistent with the observation that in Glanville fritillary butterflies, negative DDD is inferred when densities are estimated using mark-recapture method (Kuussaari *et al.* 1996), but a positive DDD is observed when the densities are experimentally manipulated (Enfjäll & Leimar 2005). Unfortunately, the effects of pre-dispersal conditions on the manifestation of DDD remain relatively poorly understood (although see Andreassen & Ims 2001; Betini *et al.* 2015).

Pre-dispersal conditions could also affect males and females differently, which might result in a differential impact on their dispersal traits. It is already known that the patterns of DDD are often asymmetric between males and females (e.g., Albrectsen & Nachman 2001; Lutz *et al.* 2015). Therefore, depending on the environmental conditions, the inherent differences in the DDD response of the two sexes could interact with the pre-dispersal conditions to yield a stronger or weaker overall DDD in the population. This could even result in the emergence of sex-biased dispersal due to differential DDD, something that has been acknowledged (Trochet *et al.* 2016) but never empirically demonstrated. Finally, because dispersal is affected by local sex ratio (Trochet *et al.* 2013), population composition in terms of number of members of the opposite sex could potentially modulate the patterns of both DDD and sex-biased dispersal.

Here we investigate some of these issues related to the robustness of DDD patterns using laboratory populations of *Drosophila melanogaster* under controlled environmental conditions. Specifically, we asked whether DDD is affected by a) pre-dispersal adult density, b) sex of the dispersers, and c) presence of mates. We found that both pre-dispersal population density and interaction between the sexes greatly affect DDD. In addition, the direction of sex-biased dispersal is shown to undergo a significant and complete reversal, via an interaction between the pre-dispersal and dispersal densities. We discuss potential reasons for the observed patterns and show how these results can impact several ecological and evolutionary processes.

## 2 MATERIAL AND METHODS

### 2.1 Fly population

All flies used in this study were collected from a large, outbred laboratory population of *D. melanogaster* (DB 4) whose detailed maintenance regime can be found elsewhere (Sah *et al.* 2013).

### 2.2 Dispersal setup and assay

Following an earlier protocol (Tung *et al.* 2017), we used a two-patch *source-path-destination* setup to study dispersal (Fig. S1). The source was a 100-mL conical flask which led into a 2-m long, transparent plastic tube (inner diameter ~1 cm) that served as the path. The other end of the path opened into a 250-mL plastic bottle that served as the destination. This end of the path protruded ~ 3 cm into the destination which prevented the flies that had reached the destination from going back into the path. The flies were introduced into the source and allowed to disperse for 4 hours. The bottle that served as the destination was replaced with another empty bottle every 30 minutes during these 4 hours, with minimum possible disturbance to the rest of the setup. The number and sex of dispersers at each of these 30-minute intervals were recorded.

### 2.3 Culturing flies for the experiments

We performed three separate experiments to investigate density dependence and sex differences in dispersal. We used four dispersal densities, namely 60, 120, 240 and 480 individuals per source container.

To eliminate any confounding effects of habitat quality, the entire dispersal setup was devoid of food and moisture, with the number of individuals in the source being the only difference across treatments during the dispersal assay. It is well known that age (Hastings 1992), kin competition (Gandon 1999) and inbreeding avoidance (Charlesworth & Charlesworth 1987) can affect the patterns of dispersal. We avoided these confounding factors by appropriate rearing of the flies as mentioned in Appendix S1 in Supporting Information.

### 2.4 Three DDD Experiments

All experiments were performed on 12-day (from egg-lay) old flies. All experiments used the same four dispersal density treatments, namely 60, 120, 240 and 480.

### Experiment 1: Mixed-sex dispersal with variable pre-dispersal densities across treatments

One day before the dispersal assay (i.e. on day 11) the adult flies were separated by sex under light CO_2_ anesthesia. The adults were then randomly assigned to the four dispersal density treatments (60, 120, 240 and 480) with a strict 1:1 sex ratio, i.e. the density treatment of 60 consisted of 30 males and 30 females, and so on. These flies were then transferred into separate 100 mL conical glass flasks (identical to the source in dispersal setup) containing ~35 mL of banana-jaggery medium. After 21 hours, the dispersal behavior of these flies were measured as described in section 2.22 (Fig. S2A). This experiment happened over 9 consecutive days with a fresh set of flies on each day, thus yielding 9 independent replicates per dispersal density. One replicate of each dispersal density was assayed on each day and a total of 8,100 flies (9 days × 900 flies/day) were used in this experiment. Flies of both sexes were present in a given dispersal setup, and the pre-dispersal density (i.e., the density during anesthesia-recovery period) was different for the four treatments.

### Experiment 2: Mixed-sex dispersal with a uniform pre-dispersal density across treatments

This experiment was identical to Experiment 1 in all respects, except for the fact that the pre-dispersal density for all the flies was held equal. For this, at the time of separation by sex under CO_2_ anesthesia, the flies were randomly assigned to 15 groups of 60 individuals with 1:1 sex ratio. Each set of 60 flies was transferred into a 35 mL plastic vial, containing ~3 mL of banana-jaggery medium. 21 hours later, at the time of dispersal setup, flies from an appropriate number of vials were mixed together to yield the density treatments (i.e., 1×60, 2×60, 4×60 and 8×60) and transferred into the corresponding source flasks (Fig. S2B). Thus, in this experiment, both sexes were assayed together, and a uniform pre-dispersal density was maintained across the four dispersal treatments.

### Experiment 3: Uni-sex dispersal with a uniform pre-dispersal density across treatments

This experiment was identical to experiment 2 in all respects, except that the males and females dispersed in the absence of the opposite sex and there were 10 replicates per density per sex. In other words, the males and females were introduced into separate dispersal setups for all four density treatments (Fig. S2C) and a total of 18,000 flies (2 sexes × 10 days × 900 flies/day/sex = 18,000 flies) were used to obtain the data. Thus, in this experiment, the two sexes were assayed independently, and the pre-dispersal density was uniform across the treatments.

Thus, over the three experiments, a total of 34,200 flies were scored for their dispersal behavior.

### 2.5 Dispersal traits

For each experiment, the obtained data included the number and sex of flies that reached the destination during each of the 30-minute intervals till the end of dispersal assay (4 hours). We also counted the flies that emigrated from the source but did not reach the destination, i.e. the flies found within the path tube at the end of dispersal assay.

Using this data, we estimated two dispersal traits as follows:

a. dispersal propensity, i.e. the proportion of flies that initiated dispersal from the source (Friedenberg 2003)

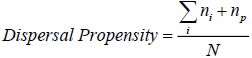
b. dispersal speed, i.e. the average speed at which the dispersers completed source-to-destination dispersal

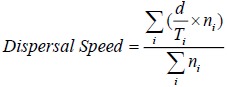

where *d* is the path distance (here, 2 m), *N* is the total number of flies introduced in the setup, *n*_*i*_ is the number of flies that reached the destination during the *t*^th^ time interval, *n*_*p*_ is the number of flies found within the path at the end of dispersal assay, and *T*_*i*_ is the time in hours since setup, at the end of *i*^th^ interval. Since the dispersal assay lasted 4 hours, with 30-minute time intervals, *T*_*i*_ ϵ (0.5, 1, 1.5, 2, 2.5, 3, 3.5, 4).

Note that dispersal propensity and speed can be independent of each other and we considered both for characterizing the dispersal behavior of our flies. Moreover, we also generated the overall temporal distribution of dispersers reaching the destination. For each dispersal setup, we divided the number of dispersers that reached the destination in each 30-minute interval by the final number of successful dispersers. Thus we obtained a distribution of dispersers against time, similar to dispersal kernels (Nathan *et al.* 2012) but with time instead of distance/location on the x-axis.

### 2.6 Statistical analyses

As stated above, each experiment was performed over multiple consecutive days, with a fresh set of age-matched flies every day. One replicate each for the four dispersal densities was assayed on each day. Thus, *day* was included in the analyses as a random blocking factor, to account for any day-specific micro-environmental variations. Data from each experiment were analyzed using a three-way mixed model ANOVA, with density (60, 120, 240 and 480) and sex (male and female) as the fixed factors, and day (1-9 for the first two experiments and 1-10 for the third experiment) as a random factor. As the propensity values are fractions, they were arcsine-square root transformed prior to the ANOVA (Zar 1999). Whenever a significant main effect was obtained, the pairwise differences were analyzed using Tukey’s HSD test. All the ANOVAs were performed using STATISTICA^®^ v5 (StatSoft. Inc., Tulsa, Oklahoma).

## 3 RESULTS

### 3.1 Pre-dispersal density modulates the effect of dispersal densities on dispersal propensity and speed

When the pre-dispersal densities were variable across the treatments (i.e., equal to the corresponding dispersal densities; Experiment 1), dispersal propensity showed a significant negative DDD (F_3,24_ = 13.66, p = 2.11 × 10^−5^; Fig. 1A). However, when the pre-dispersal density of the flies was kept uniform across all dispersal densities (Experiment 2), there was no significant effect of the latter on dispersal propensity (F_3,24_ = 1.69, p = 0.19; Fig. 1B). The results for the pairwise differences with Tukey’s p values and the associated effect sizes are given as Table S1.

**Fig. 1.**
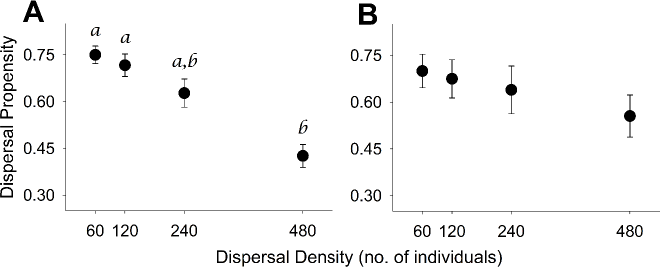
Density dependence in dispersal propensity. (**A**) Mean proportion (± SE) of the individuals dispersed vs. dispersal density for Experiment 1 (i.e., mixed-sex dispersal with variable pre-dispersal densities across treatments). (**B**) Mean proportion (± SE) of the individuals dispersed vs. dispersal density for Experiment 2 (i.e., mixed-sex dispersal with a uniform pre-dispersal density across treatments). Means with different lowercase alphabets are significantly different from each other (p<0.05 in Tukey’s HSD test). For example, in Fig. 1A, the propensity at density of 480 is significantly different from propensity at 60 and 120, whereas none of the other pairwise differences was significant. Refer to Table S1 for the exact Tukey’s p values and the associated effect sizes.

Similarly, when the pre-dispersal densities were variable (i.e. Experiment 1), there was a significant main effect of dispersal density on dispersal speed (F_3,24_ = 13.14, p = 2.81 × 10^−5^), resulting in a negative DDD (Fig. 2A). However, when the pre-dispersal densities were held uniform, even though there was a significant main effect of dispersal density (F_3,24_ = 4.78, p = 0.01), Tukey’s HSD test revealed that none of the pairwise differences were significant (Fig. 2B; see Table S2 for the exact Tukey’s p values). Thus, we show that pre-dispersal density (variable vs. uniform) can affect DDD in the context of multiple dispersal attributes.

**Fig. 2.**
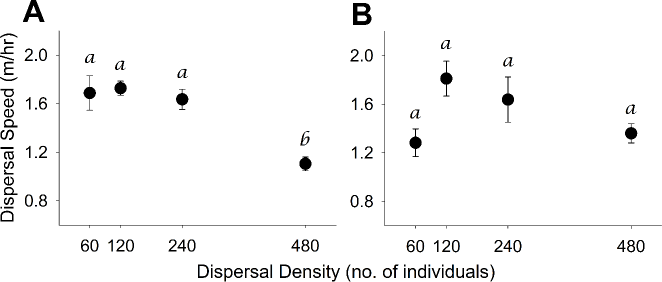
Density dependence in dispersal speed. (**A**) Mean dispersal speed (± SE) of the dispersers vs. dispersal density for Experiment 1 (i.e., mixed-sex dispersal with variable pre-dispersal densities across treatments). (**B**) Mean dispersal speed (± SE) of the dispersers vs. dispersal density for Experiment 2 (i.e., mixed-sex dispersal with a uniform pre-dispersal density across treatments). Means with different lowercase alphabets are significantly different from each other (Tukey’s HSD). Refer to Table S2 for the exact Tukey’s p values and the associated effect sizes.

### 3.2 Patterns of sex differences in DDD are modulated by pre-dispersal density

There was a significant *dispersal density × sex* interaction, in both Experiment 1 (variable pre-dispersal densities) (F_3,24_ = 10.86, p = 0.0001; Fig. 3A) and Experiment 2 (uniform pre-dispersal densities) (F_3,24_ = 5.73, p = 0.004; Fig. 3B). Two observations can be made from a comparison of these two datasets (i.e. Figs. 3A and 3B). First, the greater number of statistically significant pairwise differences for a given sex in Fig. 3A than in Fig. 3B suggests that the DDD in both sexes is amplified by variable pre-dispersal densities. Second, females have a stronger negative DDD than the males, which is evident from the greater number of significant pairwise differences across a wider range of dispersal densities for females in both Figs. 3A and 3B (see Table S3 for the exact Tukey’s p values and associated effect sizes). In other words, while the stronger DDD in Experiment 1 (variable pre-dispersal densities) than in Experiment 2 (uniform pre-dispersal densities) was reflected in both sexes, the females always exhibited a more negative density-dependent response than the males. This suggests that there are sex differences in DDD in *D. melanogaster.*

**Fig. 3.**
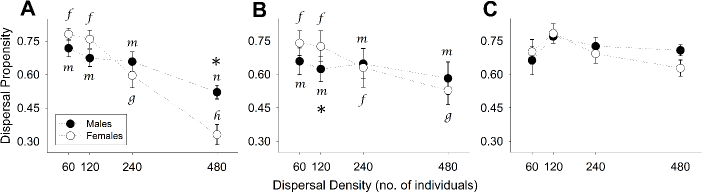
Sex differences in density dependence of dispersal propensity. (**A**) Mean dispersal propensity (± SE) of males (filled circles) and females (open circles) vs. dispersal density in Experiment 1 (i.e., mixed-sex dispersal with variable pre-dispersal densities across treatments). (**B**) Mean dispersal propensity (± SE) of males (filled circles) and females (open circles) vs. dispersal density in Experiment 2 (i.e., mixed-sex dispersal with a uniform pre-dispersal density across treatments). (**C**) Mean dispersal propensity (± SE) of males (filled circles) and females (open circles) vs. dispersal density in Experiment 3 (i.e., uni-sex dispersal with a uniform pre-dispersal density across treatments). For significant *dispersal density × sex* interactions, two kinds of pairwise comparisons are performed (using Tukey’s HSD). First, for a given sex, the extent of density dependence is examined by comparing the propensity means across the four dispersal densities (significant differences denoted using different lower-case alphabets; see Table S3 for the exact Tukey’s p values and the associated effect sizes). Second, for each dispersal density, the male and female dispersal propensities were compared (significant differences denoted by asterisks; see Table S4 for the exact Tukey’s p values and the associated effect sizes).

The difference in the strength of DDD between males and females affected the sex bias in dispersal propensity. In Experiment 1, significant sex-biased dispersal was seen only at the highest density (i.e., 480 individuals), with higher male dispersal than female dispersal (Fig. 3A). However, in Experiment 2, significant sex-biased dispersal was observed only at an intermediate density (i.e., 120 individuals), where a female-biased dispersal was observed (Fig. 3B; see Table S4 for the exact Tukey’s p values and associated effect sizes). In other words, pre-dispersal density interacted with dispersal density to determine the direction of sex-biased dispersal.

### 3.3 Patterns of sex-biased dispersal are produced by interaction between the sexes

When males and females were assayed separately while maintaining a uniform pre-dispersal density (Experiment 3), the *dispersal density × sex* interaction was not significant (F_3,27_ = 0.54, p = 0.66; Fig. 3C). In other words, males and females showed different dispersal propensities only when they dispersed together (Figs, 3A-3B) and not when they dispersed on their own (Fig. 3C). Hence, we concluded that in *D. melanogaster,* the presence of both sexes is a necessary condition for sex-biased dispersal to happen, at least under the given densities and environmental conditions.

Another interesting observation was that, in Experiment 1, there was a clear difference in the temporal patterns of male and female dispersal among the density treatments. For the proportion of males completing dispersal per time interval, there were no observable differences across the treatment densities (Fig. 4A). In contrast, the females showed a striking difference between the highest density (i.e., 480) and other treatments. While the temporal pattern of dispersal for females was quite consistent for the three lower densities (i.e., 60, 120 and 240), with a single, distinct peak at 1 hour, the distribution changed completely at the highest density (i.e., 480) (Fig. 4B). At densities lower than 480, the majority (>50%) of female dispersers completed dispersal within the first 1.5 hours, whereas at 480 individuals, the dispersal was considerably delayed.

**Fig. 4.**
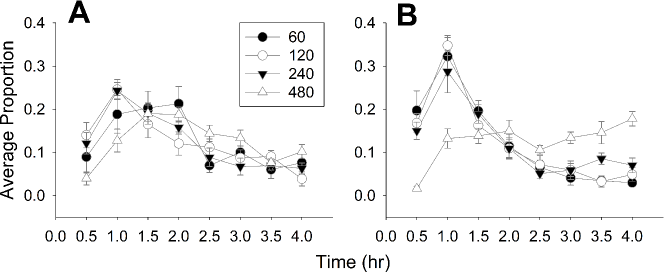
Temporal distribution of dispersers in Experiment 1 (i.e., mixed-sex dispersal with variable pre-dispersal densities across treatments). (**A**) Average proportion (± SE) of males reaching the destination vs. time. (**B**) Average proportion (± SE) of females reaching the destination vs. time.

## 4 DISCUSSION

### 4.1 Density dependence of both dispersal propensity and dispersal speed is affected by the pre-dispersal density

A strong negative density-dependence was seen in dispersal propensity when the flies were maintained at pre-dispersal densities equal to the respective dispersal densities (Fig. 1A), but not when all the flies experienced a constant pre-dispersal density (Fig. 1B). Similar results were obtained for dispersal speed as well (cf. Figs. 2A and 2B), as none of the pairwise differences among the dispersal densities was significant in Experiment 2 (Fig. 2B). Thus, even in an otherwise constant environment, something as transient as the density faced by the individuals for a brief period prior to the dispersal event can determine whether DDD is exhibited or not.

In the dispersal literature, negative DDD in asocial and non-territorial organisms has primarily been explained in terms of availability of food (Lambin 1994) and availability of mates (Kokko & Rankin 2006). However, similar larval rearing conditions and the absence of food and moisture in the dispersal setup ensured that there were no differences in terms of resources across the dispersal density treatments. The maintenance of a strict 1:1 ratio in Experiments 1 and 2 ensured that in these two experiments, there was no difference in terms of number of individuals of the opposite sex either. Thus, we explicitly controlled for the two most widely cited reasons for negative DDD. That is why our result is rather unexpected, and to the best of our knowledge, this is the first report of differences in adult pre-dispersal density leading to negative DDD.

One reason for the observed negative density-dependence of dispersal propensity and speed could be the detrimental effects of adult crowding. In *D. melanogaster,* even brief periods of enhanced adult crowding can reduce female fecundity (Joshi *et al.* 1998) and adult longevity (Joshi & Mueller 1997), which suggests a physiological change in the condition of individuals. Thus, with increasing pre-dispersal densities, the flies could have been under increased stress, which could lead to a decrease in the dispersal propensity and speed (Figs. 1A and 2A), and ultimately manifest as negative DDD (however, see section 4.3).

### 4.2 Sex differences in DDD and reversal in the direction of sex-biased dispersal

Under both variable and uniform pre-dispersal densities (Experiments 1 and 2, respectively), the *dispersal density × sex* interaction was significant for dispersal propensity (Figs. 3A and 3B). In both the experiments, the females showed a stronger negative DDD (as seen in a greater number of pairwise differences in Figs. 3A and 3B) than the males. Sex-specific differences in DDD are well known in the dispersal literature (Albrectsen & Nachman 2001; Lutz *et al.* 2015), but to the best of our knowledge, this is the first demonstration of sex differences in DDD in *D. melanogaster.* Furthermore, there was no significant sex bias in dispersal propensity in all but one density each for Experiments 1 and 2. Interestingly though, we found significant male-biased dispersal at a density of 480 individuals in Experiment 1 (denoted by * in Fig. 3A), but significant female-biased dispersal at a density of 120 in Experiment 2 (denoted by * in Fig. 3B). Although the modulation of sex-biased dispersal due to density-dependence is possible in principle (Trochet *et al.* 2016), to our knowledge, this is the first empirical demonstration of this phenomenon and that of a complete reversal in the direction of sex-biased dispersal within the same population.

### 4.3 Asymmetric DDD in males and females is caused by an interaction between the two sexes

Comparison of the results from Experiments 2 and 3 revealed that there is a difference in DDD only when the two sexes disperse together (Fig. 3B) and not when they disperse separately (Fig. 3C). To the best of our knowledge, this is the first study that compares the DDD of males and females dispersing in presence and absence of each other. Sex-biased dispersal has been recorded in a wide variety of taxa, and a number of hypotheses have been proposed to explain its origin (reviewed in Trochet *et al.* 2016). However, the observation that sex-biased dispersal can emerge as a result of interaction between males and females at different densities, even when they have intrinsically similar dispersal patterns, is a novel one.

As discussed above (section 4.1), the negative DDD observed in Experiment 1 could be due to a change in the physiological condition of the individuals at high densities. The fact that the females showed a stronger negative DDD than the males (Figs. 3A and 3B) indicated that the females are affected more than the males. However, if that were the case, then one would also expect a significant *dispersal density × sex* interaction in Experiment 3, which was not seen (Fig. 3C). Taken together, these observations suggest that, at least for the conditions of the present study, the presence of mates was a necessary condition for the manifestation of sex differences in DDD.

The asymmetry in the negative DDD patterns of males and females is further supported by observations in Fig. 4. While dispersing in the presence of the opposite sex under variable pre-dispersal densities (i.e. experiment 1), females at the highest density show a very different temporal profile of dispersal compared to the other densities (Fig. 4B). On the other hand, the temporal profiles for the males are not affected so drastically by high densities (Fig. 4A).

In *D. melanogaster,* the effects of mate-harm by males are well documented. It has been shown that lifetime female fecundity (Fowler & Partridge 1989; Carazo *et al.* 2014), egg-production rate (Pitnick & García-González 2002) and female longevity (Cohet & David 1976; Lew & Rice 2005) get affected due to mating and re-mating. Moreover, female flies experience non-mating costs of exposure to males that affects traits such as longevity (Partridge & Fowler 1990). Thus, high adult density and the presence of males together could have reduced the female dispersal propensity in Experiment 1, leading to the observed overall negative DDD.

### 4.4 Implications

In recent years, several studies have investigated the role of DDD on population dynamics (Amarasekare 2004; Ims & Andreassen 2005), biological invasions (Travis *et al.* 2009), climate change response (Best *et al.* 2007), range expansions (Altwegg *et al.* 2013) and community assembly (French & Travis 2001). However, plasticity in the patterns of DDD itself would mean that the results and predictions from such studies might not hold across different environmental scenarios. For instance, modulation of negative DDD by pre-dispersal density, as observed in our study (Figs. 1 and 2), can lead to reduced spatial connectivity in fragmented populations, thereby reducing the chances of evolutionary rescue from extinctions (Andreassen & Ims 2001; Schiffers *et al.* 2013). Thus, this result suggests that environment-dependent heterogeneity in dispersal patterns can affect key population-level processes, and needs to be explicitly accounted for in theoretical and empirical studies (Hawkes 2009).

In addition, the results from Experiments 1, 2 and 3, on the environment-dependent asymmetry in male and female dispersal, have several key implications. First, factors like sex ratio (Pérez-González & Carranza 2009) and breeding site availability (Arlt & Pärt 2008) have been hypothesized as potential reasons for variations in sex-biased dispersal. However, to the best of our knowledge, no empirical study has unequivocally identified a cause for changing patterns of sex-biased dispersal. Our results show that even such a seemingly innocuous factor as adult density for a short period prior to dispersal, can completely alter the pattern of sex-biased dispersal. Populations under natural conditions are expected to experience much greater variations in abiotic and biotic factors, many of which could potentially affect the two sexes differentially. This implies that sex bias in dispersal is perhaps even less of a robust phenomenon than what is normally believed, and extrapolations across populations (let alone species) should be made with extreme care.

Second, sex-biased dispersal is known to shape phenomena such as invasion success (Miller *et al.* 2011; Miller & Inouye 2013) and adaptive divergence (Fraser *et al.* 2004). However, these studies typically assume a constant pattern of sex bias, which is unlikely to be true under varying natural conditions. Thus, the implications of density- or environment-dependent switching of the direction of sex-biased dispersal would be an interesting area for future investigations.

Third, at the end of dispersal, biased settlement with respect to population composition in a new habitat is expected to promote population divergence (Arendt 2015). Therefore, pre-dispersal density dependent switching between male- and female-biased dispersal could manifest as strong founder effects in newly colonized patches at range fronts, potentially affecting the ecological and evolutionary processes that follow colonization (Ibrahim *et al.* 1996). Also, in addition to these direct and immediate founder effects, population composition in newly colonized patches can have indirect and extended effects on the future dispersal events following colonization (Le Corre & Kremer 1998). In other words, while environment can influence the composition of a dispersing population, the resulting skew in the composition, in turn, can be a potential determinant of the subsequent dispersal. It is, therefore, timely to investigate the role of population composition, both as a consequence and as a cause of dispersal.

## Acknowledgements

We thank Sripad Joshi and Shrinidhi Mahishi for help with the experiments. AM and ST were supported by Junior and Senior Research Fellowships, respectively, of the Council for Scientific and Industrial Research, Government of India. VRSS was supported through the GE Foundation Scholar Leaders Program. MAS and SS were supported through the INSPIRE fellowship of the Department of Science and Technology, Government of India. This study was supported by a research grant (#EMR/2014/000476) from SERB, Department of Science and Technology, Government of India and internal funding from IISER-Pune.

## Online Supporting Information for

### Appendix S1: Detailed fly culturing methods

Two cages with ~2400 adult flies each, derived from the DB 4 population (section 2.1 in main text) were maintained. To collect eggs for a particular day of the assay, one of the two cages was supplied with live yeast plate for ~24 hours. Fresh banana-jaggery medium was then supplied to the flies, allowing the females to oviposit, for 12 hours. After this period, ~50 eggs each were collected into forty 35-mL plastic vials containing ~6 mL of banana-jaggery medium. Following this, a plate with live yeast paste was provided to the flies again, to enable another round of egg collection. The same procedure was followed for the other population cage as well, with a constant difference of 24 hours between the cages. Therefore, it was possible to collect eggs every day, alternatively from two cages of the same population, for 10 days. The large breeding population size (~2400) in each cage ensured that the flies were not inbred, and random sampling from a large number of eggs during the collection reduced the chances of kin being sampled together. Moreover, as flies from a single set of collected eggs were used for the dispersal assay on a particular day, they were all of the same age (12^th^ day from egg collection) at the time of assay.

**Fig. S1.**
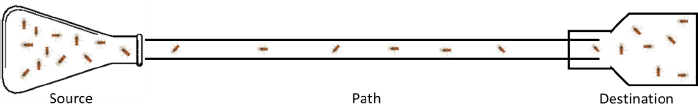
Schematic diagram of the two-patch dispersal setup. The *source* is a 100-mL glass flask (total volume ~135 mL). The *path* is a transparent plastic tube with inner diameter ~1 cm. The *destination* is a 250-mL plastic bottle. Both the source and the destination were devoid of food and moisture.

**Fig. S2.**
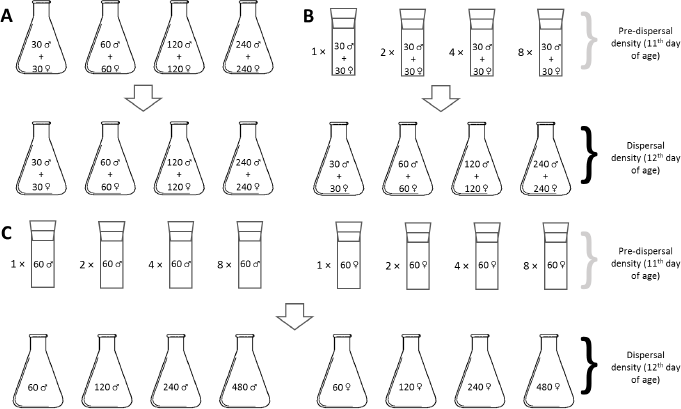
Experimental design. (**A**) Experiment 1 (4 treatments; 9 replicates per treatment) (**B**) Experiment 2 (4 treatments; 9 replicates per treatment) (**C**) Experiment 3 (8 treatments; 10 replicates per treatment)

**Table S1.** Pairwise differences for dispersal propensity in Experiment 1 (variable pre-dispersal densities) and Experiment 2 (uniform pre-dispersal density). For significant pairwise differences (Tukey’s *p* < 0.05), effect sizes (Cohen’s *d*) 95% CI are computed.

| Pre-dispersal density | Densities | Tukey's $p$ | Cohen's $d \pm 95\% \text{ CI}$ | Effect size interpretation |
| --- | --- | --- | --- | --- |
| **Variable**<br>**(Experiment 1)** | 60 – 120 | 0.96805 | - | - |
|  | 60 – 240 | 0.40071 | - | - |
| | 60 – 480 | 0.00203* | $3.306737 \pm 1.421447$ | Large |
|  | 120 – 240 | 0.66948 | - | - |
| | 120 – 480 | 0.00617* | $2.656413 \pm 1.267553$ | Large |
|  | 240 – 480 | 0.08 | - | - |
| **Uniform**<br>**(Experiment 2)** | 60 – 120 | Main effect not significant | - | - |
|  | 60 – 240 |  | - | - |
|  | 60 – 480 |  | - | - |
|  | 120 – 240 |  | - | - |
|  | 120 – 480 |  | - | - |
|  | 240 – 480 |  | - | - |

**Table S2.**
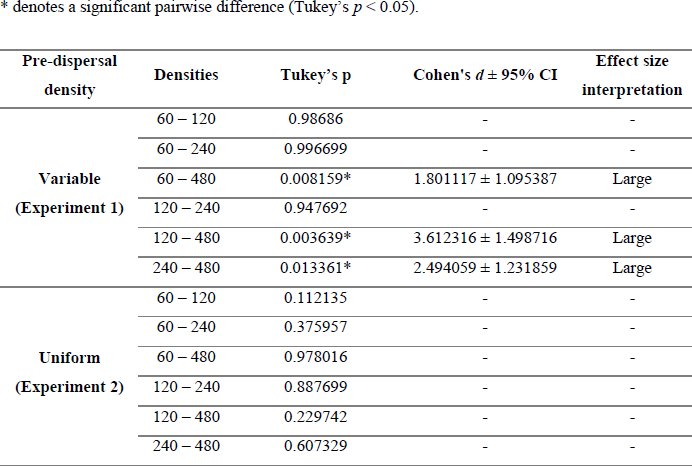
Pairwise differences for dispersal speed in Experiment 1 (variable pre-dispersal densities) and Experiment 2 (uniform pre-dispersal density). For significant pairwise differences (Tukey’s *p* < 0.05), effect sizes (Cohen’s *d*) with 95% CI are computed.

**Table S3.** Within-sex pairwise differences for dispersal propensity in Experiment 1 (variable pre-dispersal densities) and Experiment 2 (uniform pre-dispersal density). For significant pairwise differences (Tukey’s *p* < 0.05), effect sizes (Cohen’s *d*) with 95% CI are computed.

| Pre-dispersal density | Sex | Densities | Tukey's $p$ | Cohen's $d \pm 95\%$ CI | Effect size interpretation |
| --- | --- | --- | --- | --- | --- |
| **Variable**<br>**(Experiment 1)** | Males | 60 – 120 | 0.925705 | - | - |
|  |  | 60 – 240 | 0.820068 | - | - |
| | | 60 – 480 | 0.001099* | $1.903589 \pm 1.113719$ | Large |
|  |  | 120 – 240 | 0.999994 | - | - |
| | | 120 – 480 | 0.019294* | $1.490408 \pm 1.044378$ | Large |
| | | 240 – 480 | 0.035166* | $1.214002 \pm 1.005462$ | Large |
|  | Females | 60 – 120 | 0.998385 | - | - |
| | | 60 – 240 | 0.001293* | $1.431496 \pm 1.035546$ | Large |
| | | 60 – 480 | $>0.000001^*$ | $4.011698 \pm 1.603458$ | Large |
| | | 120 – 240 | 0.005694* | $1.13081 \pm 0.99506$ | Large |
| | | 120 – 480 | $>0.000001^*$ | $3.319658 \pm 1.424658$ | Large |
| | | 240 – 480 | 0.000020* | $1.762668 \pm 1.088688$ | Large |
| **Uniform**<br>**(Experiment 2)** | Males | 60 – 120 | 0.947242 | - | - |
|  |  | 60 – 240 | 0.999894 | - | - |
|  |  | 60 – 480 | 0.311063 | - | - |
|  |  | 120 – 240 | 0.995785 | - | - |
|  |  | 120 – 480 | 0.914312 | - | - |
|  |  | 240 – 480 | 0.537093 | - | - |
|  | Females | 60 – 120 | 1.000000 | - | - |
|  |  | 60 – 240 | 0.101614 | - | - |
| | | 60 – 480 | 0.000037* | $1.184309 \pm 1.001679$ | Large |
|  |  | 120 – 240 | 0.071835 | - | - |
| | | 120 – 480 | 0.000025* | $0.987034 \pm 0.978594$ | Large |
| | | 240 – 480 | 0.045248* | $0.443643 \pm 0.935253$ | Low |

**Table S4.** Sex-bias in dispersal propensity in Experiment 1 (variable pre-dispersal densities) and Experiment 2 (uniform pre-dispersal density). For significant pairwise differences (Tukey’s *p* < 0.05), effect sizes (Cohen’s *d*) with 95% CI are computed.

| Pre-dispersal density | Densities | Tukey's $p$ | Cohen's $d \pm 95\% \text{ CI}$ | Effect size interpretation |
| --- | --- | --- | --- | --- |
| **Variable**<br>**(Experiment 1)** | 60m – 60f | 0.628952 | - | - |
|  | 120m – 120f | 0.294901 | - | - |
|  | 240m – 240f | 0.739523 | - | - |
| | 480m – 480f | 0.001862* | $1.641389 \pm 1.068259$ | Large |
| **Uniform**<br>**(Experiment 2)** | 60m – 60f | 0.232469 | - | - |
| | 120m – 120f | 0.016373* | $0.538466 \pm 0.940546$ | Medium |
|  | 240m – 240f | 1.000000 | - | - |
|  | 480m – 480f | 0.820914 | - | - |

